# Genetic manipulation of an *Ixodes scapularis* cell line

**DOI:** 10.1101/2023.09.08.556855

**Authors:** Nisha Singh, Agustin Rolandelli, Anya J. O’Neal, L. Rainer Butler, Sourabh Samaddar, Hanna J. Laukaitis-Yousey, Matthew Butnaru, Stephanie E. Mohr, Norbert Perrimon, Joao H. F. Pedra

## Abstract

Although genetic manipulation is one of the hallmarks in model organisms, its applicability to non-model species has remained difficult due to our limited understanding of their fundamental biology. For instance, manipulation of a cell line originated from the blacklegged tick *Ixodes scapularis,* an arthropod that serves as a vector of several human pathogens, has yet to be established. Here, we demonstrate the successful genetic modification of the commonly used tick ISE6 line through ectopic expression and clustered regularly interspaced palindromic repeats (CRISPR)/CRISPR-associated protein 9 (Cas9) genome editing. We performed ectopic expression using nucleofection and attained CRISPR-Cas9 editing via homology dependent recombination. Targeting the E3 ubiquitin ligase X-linked inhibitor of apoptosis (*xiap*) and its substrate *p47* led to alteration in molecular signaling within the immune deficiency (IMD) network and increased infection of the rickettsial agent *Anaplasma phagocytophilum* in *I. scapularis* ISE6 cells. Collectively, our findings complement techniques for genetic engineering of ticks *in vivo* and aid in circumventing the long-life cycle of *I. scapularis,* of which limits efficient and scalable molecular genetic screens.

**Importance:** Genetic engineering in arachnids has lagged compared to insects, largely because of substantial differences in their biology. This study unveils the implementation of ectopic expression and CRISPR-Cas9 gene editing in a tick cell line. We introduced fluorescently tagged proteins in ISE6 cells and edited its genome via homology dependent recombination. We ablated the expression of *xiap* and *p47*, two signaling molecules present in the immune deficiency (IMD) pathway of *I. scapularis*. Impairment of the tick IMD pathway, an analogous network of the tumor necrosis factor receptor in mammals, led to enhanced infection of the rickettsial agent *A. phagocytophilum*. Altogether, our findings provide a critical technical resource to the scientific community to enable a deeper understanding of biological circuits in the blacklegged tick *Ixodes scapularis*.

## Introduction

The blacklegged tick *Ixodes scapularis* is a medically relevant chelicerate that transmits several bacteria, viruses and protozoa to humans and other animals (1, 2). To date, inefficient methods for genetic manipulation in *I. scapularis* makes this organism mostly intractable, which leaves significant fundamental gaps in the biology of this ectoparasite. As an example, ectopic expression is a robust tool for elucidating gene function and discovering new phenotypes. However, tick cell lines are reportedly refractory to established transfection methods (3). Additionally, the use of clustered regularly interspaced short palindromic repeats (CRISPR)/CRISPR-associated protein 9 (Cas9), the gold standard for studying functional genomics in model organisms (4–6), remains challenging despite the recent strides made in genome sequencing (7–9) and the documented *in vivo* application of CRISPR-Cas9 editing to score morphological phenotypes in *I. scapularis* (10). Currently, RNA interference (RNAi) is a widely accepted technique to study functional genomics in ticks (11) but this approach presents limitations, such as off-target effects and transient or low knockdown efficiency (12). Thus, there is a pressing need for the development of genetic tools to manipulate the biology of *I. scapularis* and better understand interactions between this arthropod vector and microbes it encounters.

In this study, we report ectopic expression and CRISPR-Cas9 gene editing of the commonly used ISE6 cell line originated from *I. scapularis*. We indicate the role of the E3 ubiquitin ligase X-linked inhibitor of apoptosis (*xiap*) and *p47* in activating the immune deficiency (IMD) pathway (13–15). The IMD network is analogous to the tumor necrosis factor (TNF) receptor pathway in mammals (16, 17) and acts as a primary defense against infection of Gram-negative bacteria in ticks (13–15). We ectopically express fluorescently tagged *xiap* and *p47* in the ISE6 cell line. We verify XIAP-p47 interactions and indicate their subcellular localization within tick cells. Importantly, CRISPR-Cas9 targeting of *xiap* led to impaired IMD pathway activation and increased infection of the rickettsial agent *Anaplasma phagocytophilum*. Taken together, these studies will aid in circumventing *in vivo* genetic methodologies that are restricted to the two-year lifespan of *I. scapularis* (2, 18). Our findings will also pave the way for the development of streamlined and scalable screening strategies to investigate cell biology and molecular mechanisms in ticks.

## Results

### Ectopic expression and XIAP-p47 interactions in a tick cell line

Labeling of organelles within cells remains an important tool in cell biology to visualize proteins of interest and identify their molecular interactions (19, 20). To determine whether fluorescent probes might be used to visualize subcellular structures in *I. scapularis*, we stained organelles within the ISE6 cell line through commonly used molecular dyes for plasma membrane, lysosome, mitochondria, endoplasmic reticulum, and Golgi apparatus (Figure 1A). The successful staining of these compartments in tick cells enabled the characterization of two previously identified proteins from *I. scapularis*: XIAP and p47 (13–15, 21). p47 is an enzymatic substrate of the E3 ubiquitin ligase XIAP and activates the tick IMD pathway through Kenny (also known as IKKγ/NEMO) in response to infection with the intracellular bacterium *A. phagocytophilum* or the Lyme disease spirochete *Borrelia burgdorferi* (13). The impairment of *p47* expression through RNAi in *I. scapularis* reduces Kenny accumulation, lessens phosphorylation of IKKβ (IRD5), and diminishes cleavage of the nuclear factor (NF)-κB molecule Relish in *I. scapularis* (13).

**Figure 1.**
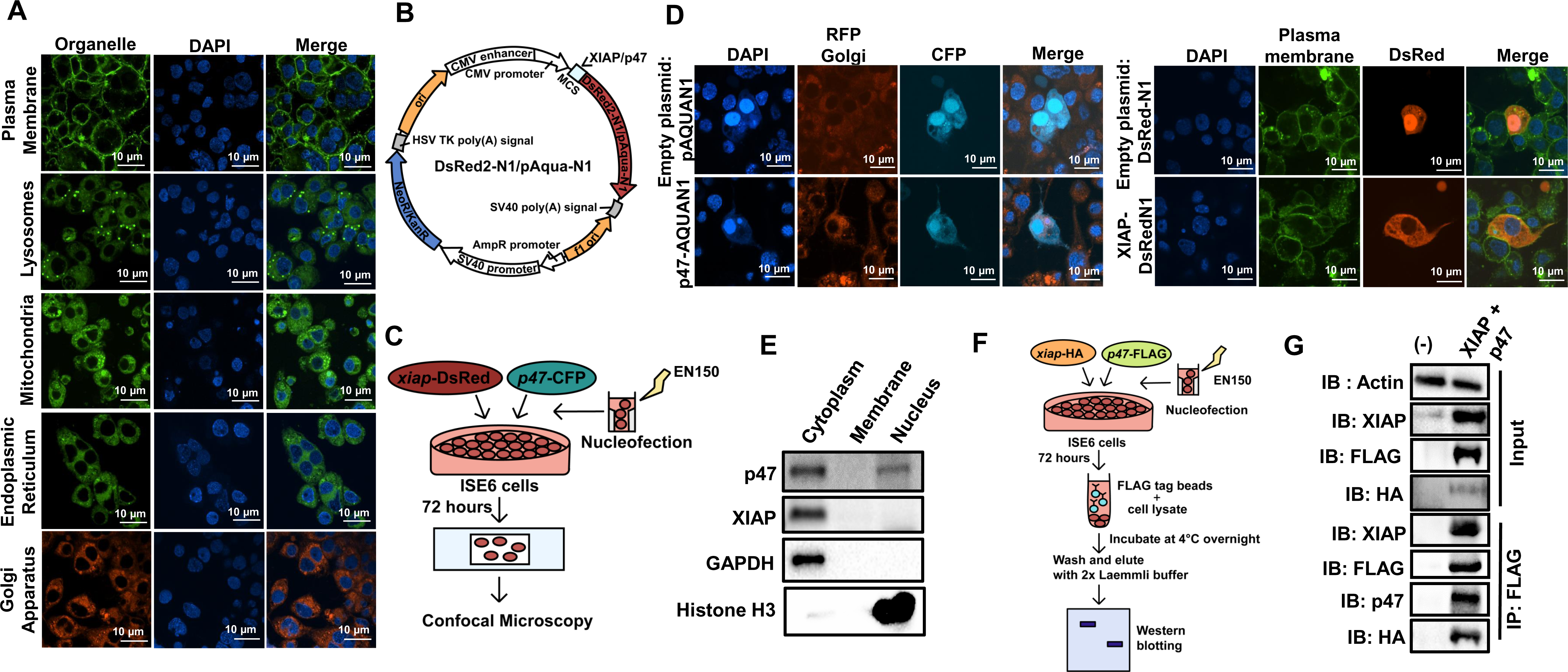
Ectopic expression in the ISE6 cell line. (A) Confocal images of ISE6 cells stained with different molecular dyes. (B) Cartoon depicting plasmids used for confocal microscopy. *Discosoma* red-N1 (DsRed2-N1) contains the fluorescent gene *DsRed* whereas p-Aquamarine-N1 (pAqua-N1) carries the fluorescent gene *Aqua*. (C) Schematic representation of ectopic expression in tick cells. (D) Ectopic expression of Aqua-tagged p47 and DsRed-tagged XIAP in ISE6 cells. ISE6 cells were nucleofected with the plasmid containing *p47*-*Aqua*, *xiap*-*DsRed* or the empty vector (DsRed-N1 or pAqua-N1). Golgi (red), plasma membrane (green) and DAPI (blue). (E) Sub-cellular fractionation of ISE6 cells. Glyceraldehyde 3-phosphate dehydrogenase (GAPDH) and histone H3 were used as cytosolic and nuclear markers, respectively. (F) Schematic representation of pull-down in ISE6 cells. *xiap*-HA and *p47*-FLAG constructs were nucleofected in ISE6 cells (3 x 10^7^). Co-transfected cells were harvested after 72 hours, and 10 mg of the lysate was incubated with the 50 µl of FLAG beads for 18 hours at 4°C. (G) The complex was immunoprecipitated (IP) using the 3X-FLAG peptide and subjected to immunoblotting (IB). Data represents one of two independent experiments.

We cloned *p47* into the pAquamarine-N1 (*p47*-Aqua) plasmid and *xiap* in the *Discosoma* red-N1 (XIAP-DsRed) plasmid (Figure 1B). Next, we successfully developed a protocol to nucleofect these plasmids into tick cells (Figure 1C). Confocal microscopy revealed that the recombinant protein p47-Aqua localized in the nucleus and the cytosol whereas XIAP-DsRed was predominantly detected in the cytosol of ISE6 cells (Figure 1D). Sub-cellular fractionation of nucleofected cells independently confirmed the location of p47 and XIAP in ISE6 cells (Figure 1E). Previously, McClure et al., 2019 demonstrated that XIAP binds and ubiquitinylates p47 in a lysine (K)-63 dependent manner in human embryonic kidney (HEK) 293T cells (13). To take advantage of the newly developed protocol for ectopic expression in tick cells, we next performed co-immunoprecipitation using the XIAP-HA and p47-FLAG vectors (Figure 1F). As shown for HEK293T cells, we detected molecular interactions between XIAP and p47 via affinity purification in ISE6 cells (Figure 1G). Taken together, these data demonstrate the ectopic expression of immune molecules in ISE6 cells and confirm protein-protein interactions in *I. scapularis*.

### CRISPR-Cas9 genome editing in the ISE6 cell line

Next, we performed *xiap* and *p47* genome editing via CRISPR-Cas9 in the ISE6 cell line. We isolated genomic DNA from ISE6 cells and designed PCR primers to amplify 1000 base pairs (bp) of the target regions. Specifically, two single guide RNAs (sgRNAs) per gene were made for *xiap* and *p47* (Supplementary Table 2). We developed a methodology for CRISPR-Cas9 through homology directed repair (HDR) (Figures 2-3). Genome manipulation via HDR displays a precise editing mechanism when a template is introduced into cells as a donor for homologous recombination (22, 23). To accomplish this, we used a cassette with an antibiotic marker and a reporter gene, along with DNA fragments homologous to the target gene. These homologous regions were positioned on either side of the donor cassette to facilitate recombination (Figures 2A and 3A). Due to the low efficiency associated with transfecting tick lines, we employed a ribonucleoprotein (RNP) delivery method to introduce the Cas9 endonuclease protein and sgRNAs in ISE6 cells (Figure S1). Following nucleofection, cells were split and selected with puromycin (Figure S2). We chose antibiotic selection over cell sorting because it remains technically unfeasible to culture tick cells from a single clone.

**Figure 2.**
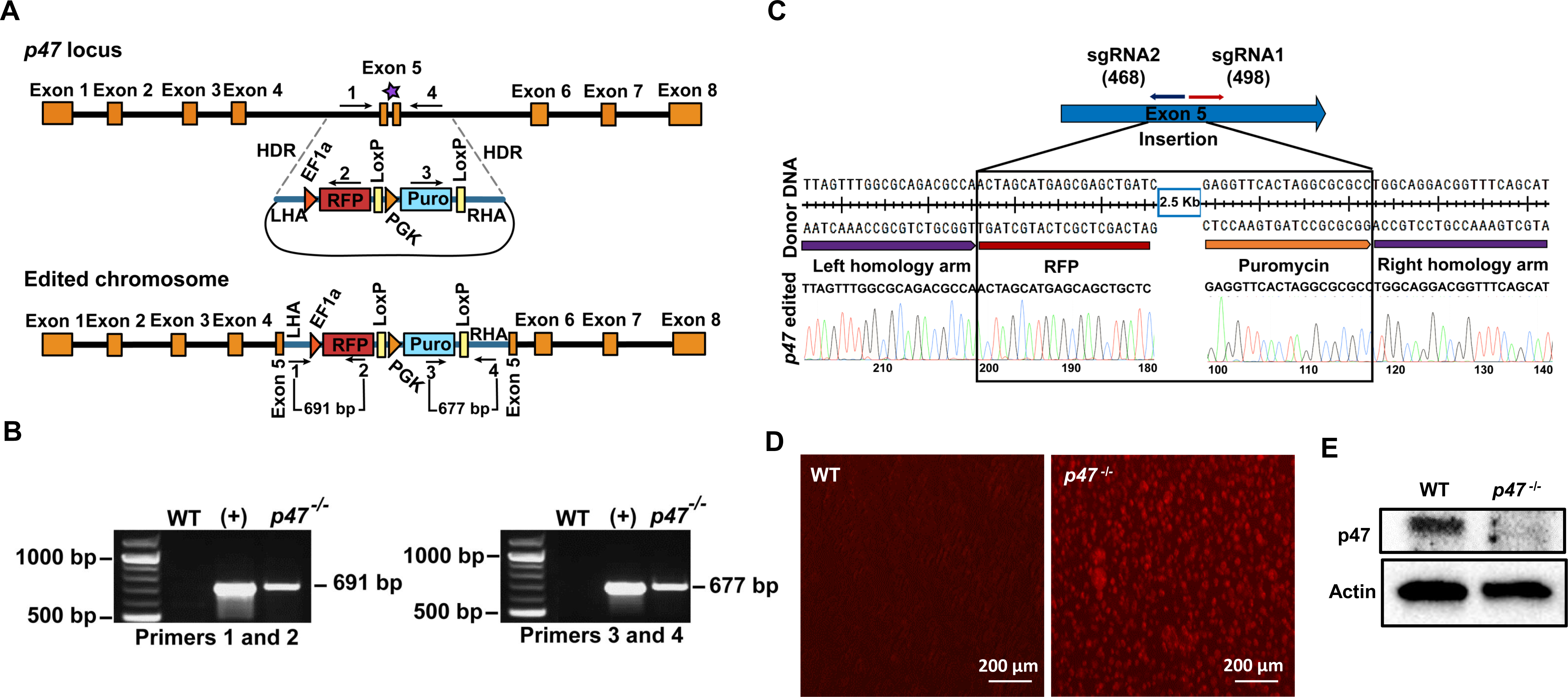
CRISPR-Cas9 *p47* editing in ISE6 cells. (A) Schematic representation of the *p47* locus and the donor construct. The orange boxes represent the eight exons of the *p47* locus, and the black line represents the intron sequence (top). The purple star on exon 5 represents the sgRNA binding and Cas9 cleavage site. The donor DNA (bottom) carries the promoter and coding sequence for the red fluorescent protein (RFP)-puromycin cassette (RFP-Puro) with the loxp (locus of X-over P1) sites in the pUC19 backbone. The donor DNA construct also carries DNA fragments of ∼600 bp in length, homologous to the *p47* gene locus, flanking the Cas9 cleavage site on the 5’ and 3’ ends for homology-directed repair (HDR). The arrows with numbers 1-4 represent primers for gene amplification analysis. (B) PCR amplification confirming the integration of the donor cassette (RFP-Puro) in the genomic DNA prepared from *p47* edited (*p47^-/-^*) cells. PCR was performed using the primer pairs 1 and 2 and 3 and 4, as mentioned in (A). The genomic DNA prepared from wildtype (WT) ISE6 cells did not show any amplification. The donor DNA was used as a positive control (+). The length in nucleotides of the amplified fragments is represented in numbers. (C) Editing was further confirmed through Sanger sequencing where the blue arrow is the targeted exon (Exon 5), red and dark blue arrows represent the sgRNA target sites with arrowheads showing the binding direction. The number in parenthesis is the binding location of sgRNA on the exon. The results were validated through fluorescence microscopy (D) and western blotting (E).

**Figure 3.**
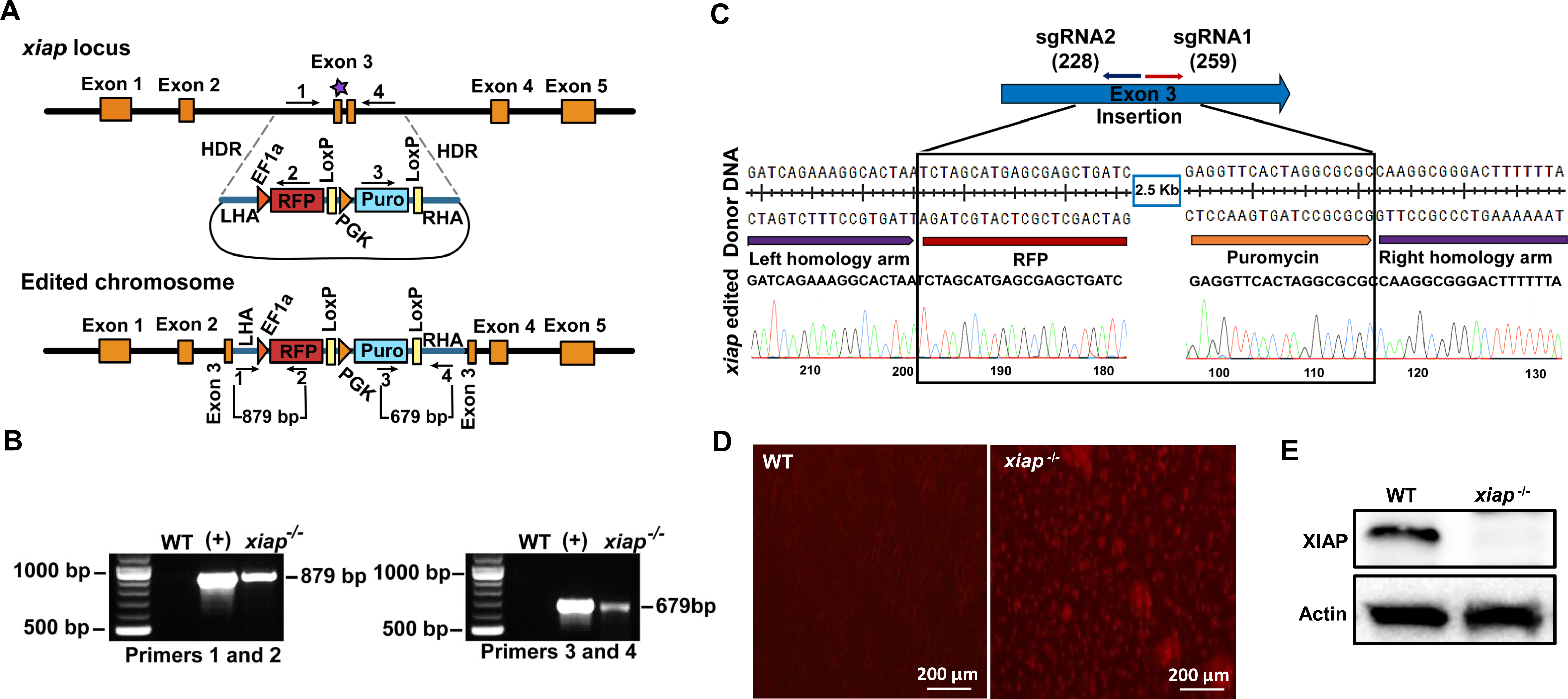
CRISPR-Cas9 *xiap* editing in ISE6 cells. (A) Schematic representation of the *xiap* locus and the donor construct. The orange boxes represent the five exons of *xiap* with sgRNA binding and the Cas9 cleavage site on exon 3 (purple star). The donor construct contains the red fluorescent protein (RFP) and puromycin cassette (RFP-Puro) along with the DNA fragments of ∼600 bp in length, homologous to the *xiap* locus, flanking the Cas9 cleavage site on the 5’ and 3’ ends for homology-directed repair (HDR). The arrows with numbers 1 and 4 represent the primers for the gene amplification analysis. (B) PCR amplification analysis confirming the integration of the donor cassette (RFP-Puro) in the genomic DNA prepared from *xiap* edited cells (*xiap^-/-^*). PCR was performed using the primer pairs 1 and 2 and 3 and 4, as mentioned in (A). The genomic DNA prepared from wildtype (WT) ISE6 cells did not show any amplification. The donor DNA was used as the positive control (+). The length in nucleotides of the amplified fragments is represented in numbers. (C) Editing was confirmed through Sanger sequencing where the blue arrow is the targeted exon (Exon 3), red and dark blue arrows indicate the sgRNA target sites with arrowheads showing the binding direction. The number in parenthesis is the binding location of sgRNA on the exon. Results were validated via fluorescence microscopy (D) and western blotting (E).

For validation of the editing event, primer sets were designed to target both *p47* and *xiap* edited cells amplifying the left homology arm and the RFP gene. Additionally, the primer pairs were designed to cover the puromycin cassette and the right homology arm (Figure 2A and 3A). PCR amplification detected the donor cassette insertion in the edited *p47^-/-^* and *xiap^-/-^* ISE6 cells (Figure 2B and 3B). Knock-in events were orthogonally confirmed through Sanger sequencing and confocal microscopy (Figures 2C-D and Figures 3C-D). Finally, western blot revealed the absence of the wildtype protein bands for p47 and XIAP in edited ISE6 cells (Figures 2E and 3E). Notably, edited *p47^-/-^*cells did not survive the puromycin selection procedure, likely due to the involvement of *p47* in growth. Disruption of a p47 homolog in the budding yeast *Saccharomyces cerevisiae*, named Shp1, leads to lethality (24, 25). Collectively, these findings confirmed the delivery of Cas9 RNPs and ablation of either *p47* and *xiap* in ISE6 cells.

### Functional disruption of *xiap* impairs IMD signaling pathway in *I. scapularis*

Given that we genetically edited *xiap* in *I. scapularis*, we then asked whether cells deficient in *xiap* (*xiap*^-/-^) were impaired for immune signaling pathways (Figure 4A) (13–15). Editing of *xiap* affected the transcription and translation of *p47*, *relish and kenny* (IKKγ/NEMO) and the cleavage of the NF-κB molecule Relish in ISE6 cells (Figures 4B-C). Next, we determined the effect of *xiap* editing in ISE6 cells during microbial stimulation with the Lyme disease spirochete *B. burgdorferi* or the rickettsial agent *A. phagocytophilum*. A significant decrease in the accumulation of Kenny and reduced nuclear translocation of N-Rel were observed in *xiap*^-/-^ cells during microbial stimulation (Figures 4D-G), indicating that functional disruption of *xiap* results in the impairment of the IMD pathway in ticks. Importantly, wild-type (WT) and *xiap^-/-^* ISE6 cells were infected with *A. phagocytophilum* for 72 hours and bacterial burden was assessed by RT-qPCR. We observed an increase in *A. phagocytophilum* load in *xiap^-/-^*ISE6 cells compared to the control treatment (Figure 4H). Taken together, these data obtained through CRISPR editing implicated the E3 ubiquitin ligase XIAP as an important molecule for the *I. scapularis* IMD pathway.

**Figure 4.**
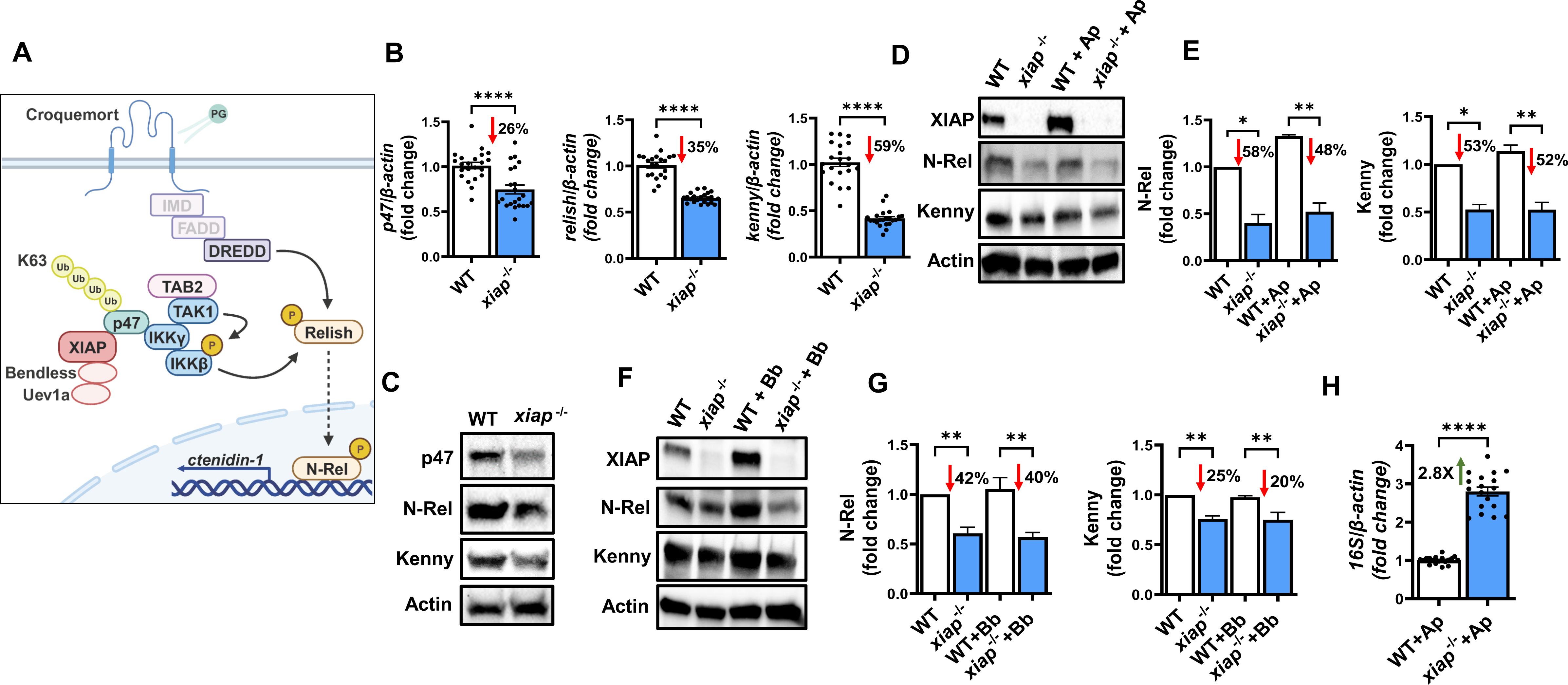
Functional disruption of *xiap* impairs the IMD signaling network in *I. scapularis*. (A) Graphical representation of the IMD pathway in ticks. (B-C) Functional disruption of *xiap* impaired the expression of molecules associated with the IMD pathway at both transcriptional (B) and translational levels (C). (D-G) 3 x 10^6^ wildtype (WT) and *xiap*^-/-^ cells were plated in a 6-well plate and stimulated with *A. phagocytophilum* (multiplicity of infection - MOI 50) or *B. burgdorferi* (MOI 50) for 15 minutes. Disruption of XIAP signaling impairs Kenny accumulation and Relish cleavage in response to (D-E) *A. phagocytophilum* infection or (F-G) *B. burgdorferi* stimulation. For data normalization, Kenny and N-Rel band densities were normalized to actin and values were divided by the uninfected WT control densitometry. Western blot images are a representative image of at least 2 independent experiments. (H) WT and *xiap*^-/-^ cells were infected with A. phagocytophilum (MOI 50). ISE6 cells were harvested after 72 hours of infection. The *A. phagocytophilum* 16S rRNA transcript was quantified by qRT-PCR and the expression data was normalized to *I. scapularis* β-actin. The qRT-PCR data show the combination of three independent experiments. Results are represented as a mean ± SEM. Statistical significance was evaluated by unpaired t test with Welch’s correction (B and H) and one way ANOVA with post-hoc Tukey (E and G). \**p*<0.05; \*\**p*<0.01; \*\*\*\**p*<0.0001.

## Discussion

Despite the recent *in vivo* application of CRISPR-Cas9 editing in *I. scapularis* (10), genetic manipulation of ticks lags behind model insects. This scientific impediment precludes a better understanding of tick biology and its interactions with microbes. Our study expands the genetic toolbox in *I. scapularis* and makes feasible ectopic expression in tick cells. For instance, we: (1) expressed two tick proteins in ISE6 cells, XIAP and p47 (13–15, 21); (2) confirmed their previous molecular interactions (13); and (3) successfully detected their subcellular location. We also developed a protocol for CRISPR-Cas9 gene editing in ISE6 cells. We provide evidence that disrupting components of the IMD pathway through a genetically edited cell line resulted in an increase of *A. phagocytophilum* burden in *I. scapularis*. Notably, our approach follows an earlier report in which Kurtti *et al*. 2008 used cationic lipid-based transfection reagents to deliver a red fluorescent protein and a selectable marker, neomycin phosphotransferase, into ISE6 cells (26). Taken together, the utilization of tick cell lines to ask biological questions offers a valuable resource for biomedical research due to its convenience and cost-effectiveness.

Recent advancements used embryo injection of CRISPR-Cas9 through the Receptor-Mediated Ovary Transduction of Cargo (ReMOT Control) (27) for direct delivery of the Cas9-RNP complex into *I. scapularis* (10). Although this technology is a breakthrough for the tick community, there are significant limitations for its applicability *in vivo*, including low survival and efficiency in addition to the long lifecycle of *I. scapularis* ticks. Our study provides a complementary approach. The technical advancements described here pave the way for exploring ancillary CRISPR-Cas9 technologies, including CRISPR activation (CRISPRa) (28) and CRISPR interference (CRISPRi) (29). Specifically, CRISPRa might be considered a valuable tool for orthogonally validating immunity and fitness studies in *I. scapularis*. By over-expressing genes of interest in *I. scapularis* cells, one may complement RNAi-based technologies and develop small- or large-scale studies, such as commonly reported genome-wide pooled CRISPRa screens in mammals and *Drosophila* (28, 30).

Collectively, the development of genetic tools for tick research offers unique avenues to identify crucial genes related to growth, physiology, immune signaling and detection of microbes. Indeed, ectopic expression and CRISPR technologies in cells might aid in the identification of antigen targets for the development of tick-based vaccines (31), epitopes associated with the α-gal allergy to red meat (32) and proteins linked to acaracide resistance (33).

## Materials and methods

### Tick cell culture

All experiments were performed under guidelines from the NIH and approved by the Institutional Biosafety Committee (IBC-00002247) at the University of Maryland, Baltimore. The embryonic cell line (ISE6) was cultured at 34°C in L15C300 medium supplemented with 10% heat inactivated fetal bovine serum (FBS, Millipore Sigma), 0.1% bovine lipoprotein concentrate (LPPC, MP Biomedicals) and 10% tryptose phosphate broth (TPP, BD). ISE6 cells were grown to confluency in T25 flasks (Greiner) and verified by PCR to be *Mycoplasma* free (Southern Biotech).

### Plasmids

*xiap* and *p47* were cloned in the pCMV-HA and pCMV-FLAG vectors, respectively, as previously described (13). For confocal microscopy, *xiap* and *p47* were cloned in DsRed2-N1(a gift from Michael Davidson, Addgene plasmid #54493) and pAQUA-N1vectors (a gift from Fabienne Merola, Addgene plasmid # 42888). Genes of interest were amplified from ISE6 cells complementary DNA (cDNA) using Phusion polymerase (NEB Biolabs). *xiap* and *p47* were cloned between the restriction sites *SacI*/*EcoRI* and *SalI*/*BamHI*, respectively. The *xiap*- and *p47*-donor DNA used in CRISPR experiments were procured from Origene technologies with a customized RFP-Puro cassette. All constructs were verified through Sanger sequencing.

### Bacteria

*Escherichia coli* BL21 (DE3) was cultured overnight at 37°C in lysogeny broth (LB) supplemented with 100 µg/ml ampicillin. *A. phagocytophilum* strain HZ was cultured in the human leukemia cell line, HL-60 cell line (ATCC, CCL-240) (34). Host-free *A. phagocytophilum* was obtained by collecting the infected-HL-60 cells at 3,260 x g for 10 minutes. The pellet was resuspended in L15C300 medium and lysed by passing through a 27 ½ gauge needle five times. Cell debris was separated by centrifugation at 750 x g for 5 minutes at 4°C. Host-free bacteria were enumerated using the following formula: number of infected HL-60 cells × 5 morulae/cell × 19 bacteria/morulae × 0.5 recovery rate (35). Low passage *B. burgdorferi* B31 clone MSK5 was cultured in Barnour-Stoenner-Kelly (BSK)-II medium supplemented with 6% normal rabbit serum at 37°C, as previously described (36, 37). Plasmid Profiling was performed by PCR amplification of necessary virulence plasmids (36).

### Antibody generation

The *I. scapularis* Kenny custom antibody used in this study was generated by Genscript. Rabbits were immunized three times with 0.2 mg of tick Kenny immunogen (amino acids 223-356). Animals were euthanized and the antiserum was obtained. Polyclonal antibodies were purified through affinity purification and tested for antigen specificity. The *I. scapularis* Relish monoclonal custom antibody used in this study was generated by Genscript. Mice were immunized three times with 0.2 mg of the tick N-Rel immunogen (Rel homology domain; amino acids 19-192). Animals were selected for cell fusion where parental clones were screened, and five positive clones were selected for subcloning. An appropriate parental clone was selected based on western blotting and the monoclonal antibody was purified by Protein A/G affinity column chromatography.

### Nucleofection

For nucleofection, 2 x 10^5^ ISE6 cells were pelleted by centrifugation at 100 x g for 10 minutes. The pellet was washed with 10 ml of 1 x PBS and resuspended in 20 µl of nucleofection SF buffer (Lonza Biosciences), in which 600 ng of the DsRed2-N1 plasmid was added to the suspension. The nucleofection mix was transferred to a multi-well cuvette and subjected to the EN150 pulse condition using a 4D-Nucleofector system (Lonza Biosciences). Following nucleofection, cells were incubated in the cuvette for 10 minutes at room temperature. ISE6 cells were added to pre-warmed L15C complete media in a 12-well plate and observed for fluorescence by microscopy after 72 hours (15).

### Pull-down assays

Following nucleofection of 4 x 10^8^ ISE6 cells, the pellet was resuspended in 800 µl of SF buffer. Twenty-five µg of either *p47*-FLAG or *xiap*-HA plasmids was added to the cell suspension. The cell suspension was split among eight nucleofection cuvettes (100 µl each) and pulsed using EN150. After 10 minutes post-nucleofection, tick cells were added to a T25 flask containing 5 ml of pre-warmed L15C complete medium and incubated at 34 °C with 1% CO_2_ for 72 hours. Cells were collected, washed twice with 1 x PBS, and lysed in immunoprecipitation lysis buffer (Thermo Scientific). The lysate (10 mg) was incubated with 300 μl of anti-FLAG cross-linked agarose beads overnight at 4°C. Beads were washed three times with 200 mM NaCl added to 1 x PBS and boiled for 5 minutes in 2X Laemmli buffer to elute proteins. Eluted proteins and input samples were analyzed by western blot for XIAP-HA and p47-FLAG detection.

### Confocal microscopy

For microscopy, 5 x 10^5^ ISE6 cells were nucleofected with 500 ng of plasmid (*xiap*- DsRed-N1 or *p47*-AQUA-N1). Tick cells were then plated on a glass coverslip (Corning). After 72 hours, cells were stained with the molecular dyes (Supplementary Table 1), as per the manufacturer’s protocol. ISE6 cells were fixed with 4% paraformaldehyde followed by 1 x PBS washes and the coverslip was mounted on a slide using Antifade gold mounting reagent with 4′,6-diamidino-2-phenylindole (DAPI) (Invitrogen) and observed under the Nikon W-1 Spinning Disk confocal microscope. The following laser channels were used: 561 nm (XIAP-DsRed), 488 nm (GFP, plasma membrane, lysosomes, mitochondria, and endoplasmic reticulum), 405 nm (DAPI), 456 nm (p47-AQUA) and 561 nm (RFP, Golgi apparatus).

### Subcellular fractionation

Subcellular fractionation was performed as previously described (38). Briefly, ISE6 cells were resuspended in 500 µl fractionation buffer and passed through a 27-gauge needle 10 times. After a 20-minute incubation on ice, the cells were centrifuged at 720 x g for 5 minutes. The supernatant was ultra-centrifuged at 100,000 x g for 1 hour to isolate the membrane fraction and the pellet was separated for the nuclear fraction. The supernatant was collected as the cytosol fraction. The pellet was washed in 400 µl of fractionation buffer, passed through 25-gauge needle and re-centrifuged at 100,000 x g for 45 minutes. The membrane pellet was resuspended in TBS containing 0.1 % Sodium dodecyl sulfate (SDS) for western blot detection.

### sgRNA and donor DNA design

Amplicons were validated using Sanger sequencing and aligned to the reference genome. Exons were identified using the ORF finder tool (https://www.bioinformatics.org/sms2/orf_find.html). Exon 3 for *xiap* and exon 5 for *p47* were selected. The CHOPCHOP server (https://chopchop.cbu.uib.no/) was used to identify guide RNA hits. The sgRNAs were selected based on the ∼20 bp sequences adjacent to NGG-PAM (protospacer adjacent motif) with 40-80% GC content and no off-target binding. The sgRNA scaffold contained the CRISPR RNA (crRNA) or the ∼20 bp target sequence, as well as the transactivating CRISPR RNA (tracrRNA) (39, 40). Two sgRNAs sequences were selected per gene, one targeting each strand (Supplementary Table 2) and customized from Synthego. The *sp*Cas9 protein and sgRNAs were combined *in vitro* to form the RNP complex which was then introduced into the ISE6 cells through nucleofection, along with the donor DNA.

To induce HDR, a donor DNA or DNA repair template was delivered to cells along with the sgRNA and Cas9 endonuclease (40–42). The donor DNA constructs targeting *xiap* and *p47* genes had the following features: (*i*) a red fluorescent protein (RFP) driven by the EF1α promoter; (*ii*) the puromycin gene for antibiotic based selection flanked by a phosphoglycerate kinase 1 (PGK) promoter; (*iii*) the loxp (locus of X-over P1) sites to flox out the puromycin cassette; and (*iv*) DNA fragments of ∼600 bp in length, homologous to the *xiap* or *p47* gene locus flanking the Cas9 cleavage site on the 5’ and 3’ ends. The resulting plasmid was of ∼7 Kb in length with the 2.5 Kb RFP-Puro cassette targeted for insertion at the *xiap* or the *p47* gene loci.

### CRISPR-Cas9 gene editing in tick cells

To prepare the RNP, 200 pmol of Cas9-NLS-tagRFP protein (Genaxxon bioscience) and 100 µM of sgRNAs for *p47* and *xiap* (Supplementary Tables 1-2) were mixed and incubated at room temperature for 20 minutes. The RNP complex together with 7 µg of donor DNA was nucleofected into 3 × 10^7^ ISE6 cells using EN150 and buffer SF via the 4D-Nucleofector system (Lonza Bioscience) (Figure S1). ISE6 cells were split (1:10) 3 days post-nucleofection and then cultured for 10 days. This process was repeated for 7 cycles followed by puromycin selection (4 µg/ml) to select for edited cells.

### Antibiotic selection

5 x 10^5^ ISE6 cells were plated on a 12-well plate and increasing concentrations of puromycin (0-10 µg/ml) were added to the cells after 24 hours. The medium containing the puromycin antibiotic was replaced every two days for a duration of 10 days. Cell viability was measured using trypan blue to determine the appropriate puromycin concentration (Figure S2).

### Quantitative reverse transcription polymerase chain reaction (qRT-PCR)

RNA was extracted from the cells preserved in TRIzol (Invitrogen) using the PureLink RNA Mini kit (Invitrogen) and the cDNA was synthesized with the Verso cDNA Synthesis Kit (ThermoFisher). Gene expression was measured using the primers listed in Supplementary Table 2.

### Western blotting

ISE6 cells (3×10^6^ cells per well) were plated in 6-well plates (Millipore Sigma) and stimulated accordingly. Protein lysate was prepared in Radio-immunoprecipitation assay (RIPA) buffer (Merck Millipore) with a protease inhibitor cocktail (Roche) and protein concentration was estimated using BCA assay (Thermo Scientific). Sodium dodecyl sulfate polyacrylamide gel electrophoresis (SDS-PAGE) samples were prepared by boiling equal amounts of protein in 6X Laemmli sample buffer (Alfa Aesar) containing 5% β-mercaptoethanol. Proteins were transferred onto PVDF membranes (Biorad) and membrane was blocked for 1 hour with 5% skimmed milk prepared in PBS-T. Primary antibodies were incubated overnight at 4°C in PBS-T and blots were washed four times in PBS-T. Blots were subsequently incubated with secondary antibodies for at least 1 hour at room temperature with gentle rocking. Blots were washed four times in PBS-T, incubated with enhanced chemiluminescence (ECL) substrate solution for 1 minute (Millipore), and imaged.

### Statistical Analysis

Three independent experiments were performed for each set of experiments. Means ± standard error of the mean (SEM) was plotted, and statistical significance was assessed by the unpaired *t* test with Welch’s correction or one-way analysis of variance (ANOVA). GraphPad PRISM® (GraphPad Software version 9.1.0) was used for statistical analyses and GraphPad Quickcals program was used for outlier detection (https://www.graphpad.com/quickcalcs/Grubbs1.cfm). A *p* value<0.05 was considered statistically significant.

## Acknowledgements

We acknowledge members of the Pedra laboratory for providing insightful discussions and manuscript feedback. We thank Jon Skare (Texas A&M University Health Science Center) for providing the *B. burgdorferi* B31 strain, clone MSK5; Ulrike G. Munderloh (University of Minnesota) for supplying the ISE6 cell line; Joseph Mauban (University of Maryland School of Medicine) for aiding in confocal microscopy; Biopolymer/Genomics core facility for Sanger sequencing and Francy E. Cabrera Paz (University of Maryland School of Medicine) for providing administrative support. This work was supported by grants from the National Institutes of Health (NIH) to AJO (F31AI152215), LRB (F31AI167471), HJL-Y (T32AI162579), JHFP (R01AI134696, R01AI116523 and P01AI138949), NP, SEM and JHFP (R21 AI168592). NP, SEM and JHF were also supported in-kind by the Fairbairn Family Lyme Research Initiative. NP is an investigator of the Howard Hughes Medical Institute. Figure 4A was created with BioRender.com. The content is solely the responsibility of the authors and does not represent the official views of the NIH, the Department of Health and Human Services or the United States government.

## Author contributions

NS and JHFP designed the study. NS, AR, AJO, LRB, HJLY and SS performed the experiments. NS and JHFP wrote the manuscript. LRB and HJLY helped in creating schematics. MB, SEM and NP provided intellectual insights and scientific advice. All authors analyzed the data and contributed to editing of the manuscript. JHFP supervised the study.

## Supplementary Materials

Figures S1 to S2 Tables S1 to S2

## Notes

### Competing Interest Statement

The authors have declared no competing interest.

## References

1. Eisen L. 2020. Stemming the rising tide of human-biting ticks and tickborne diseases, United States. Emerg Infect Dis 26:641–647.

2. Eisen RJ, Eisen L. 2018. The blacklegged tick, *Ixodes scapularis*: an increasing public health concern. Trends Parasitol 34:295–309.

3. O’Neal AJ, Singh N, Mendes MT, Pedra JHF. 2021. The genus *Anaplasma*: drawing back the curtain on tick-pathogen interactions. Pathog Dis 79: ftab022.

4. Zhang F, Wen Y, Guo X. 2014. CRISPR/Cas9 for genome editing: progress, implications and challenges. Hum Mol Genet 23:R40–6.

5. Gratz SJ, Rubinstein CD, Harrison MM, Wildonger J, O’Connor-Giles KM. 2015. CRISPR-Cas9 genome editing in *Drosophila*. Curr Protoc Mol Biol 111:31.2.1-31.2-20.

6. Bassett AR, Liu JL. 2014. CRISPR/Cas9 and genome editing in *Drosophila*. J Genet Genomics 41:7–19.

7. Gulia-Nuss M, Nuss AB, Meyer JM, Sonenshine DE, Roe RM, Waterhouse RM, Sattelle DB, de la Fuente J, Ribeiro JM, Megy K, Thimmapuram J, Miller JR, Walenz BP, Koren S, Hostetler JB, Thiagarajan M, Joardar VS, Hannick LI, Bidwell S, Hammond MP, Young S, Zeng Q, Abrudan JL, Almeida FC, Ayllon N, Bhide K, Bissinger BW, Bonzon-Kulichenko E, Buckingham SD, Caffrey DR, Caimano MJ, Croset V, Driscoll T, Gilbert D, Gillespie JJ, Giraldo-Calderon GI, Grabowski JM, Jiang D, Khalil SMS, Kim D, Kocan KM, Koci J, Kuhn RJ, Kurtti TJ, Lees K, Lang EG, Kennedy RC, Kwon H, Perera R, Qi Y, et al. 2016. Genomic insights into the *Ixodes scapularis* tick vector of Lyme disease. Nat Commun 7:10507.

8. Miller JR, Koren S, Dilley KA, Harkins DM, Stockwell TB, Shabman RS, Sutton GG. 2018. A draft genome sequence for the *Ixodes scapularis* cell line, ISE6. F1000Res 7:297.

9. De S, Kingan SB, Kitsou C, Portik DM, Foor SD, Frederick JC, Rana VS, Paulat NS, Ray DA, Wang Y, Glenn TC, Pal U. 2023. A high-quality *Ixodes scapularis* genome advances tick science. Nat Genet 55:301–311.

10. Sharma A. PMN, Reyes J. B., Chana R., Yim W. C., Heu C. C., et al. 2020. Cas9-mediated gene-editing in the black-legged tick, Ixodes scapularis, by embryo injection and ReMOT control. iScience 25:103781.

11. de la Fuente J, Kocan KM, Almazan C, Blouin EF. 2007. RNA interference for the study and genetic manipulation of ticks. Trends Parasitol 23:427–33.

12. Sledz CA, Williams BR. 2005. RNA interference in biology and disease. Blood 106:787–94.

13. McClure Carroll EE, Wang X, Shaw DK, O’Neal AJ, Oliva Chavez AS, Brown LJ, Boradia VM, Hammond HL, Pedra JHF. 2019. p47 licenses activation of the immune deficiency pathway in the tick *Ixodes scapularis*. Proc Natl Acad Sci U S A 116:205–210.

14. Shaw DK, Wang X, Brown LJ, Chavez AS, Reif KE, Smith AA, Scott AJ, McClure EE, Boradia VM, Hammond HL, Sundberg EJ, Snyder GA, Liu L, DePonte K, Villar M, Ueti MW, de la Fuente J, Ernst RK, Pal U, Fikrig E, Pedra JH. 2017. Infection-derived lipids elicit an immune deficiency circuit in arthropods. Nat Commun 8:14401.

15. O’Neal AJ, Singh N, Rolandelli A, Laukaitis HJ, Wang X, Shaw DK, Young BD, Narasimhan S, Dutta S, Snyder GA, Samaddar S, Marnin L, Butler LR, Mendes MT, Cabrera Paz FE, Valencia LM, Sundberg EJ, Fikrig E, Pal U, Weber DJ, Pedra JHF. 2023. Croquemort elicits activation of the immune deficiency pathway in ticks. Proc Natl Acad Sci U S A 120:e2208673120.

16. Lemaitre B, Kromer-Metzger E, Michaut L, Nicolas E, Meister M, Georgel P, Reichhart JM, Hoffmann JA. 1995. A recessive mutation, immune deficiency (*imd*), defines two distinct control pathways in the *Drosophila* host defense. Proc Natl Acad Sci USA 92:9465–9.

17. Kleino A, Silverman N. 2014. The *Drosophila* IMD pathway in the activation of the humoral immune response. Dev Comp Immunol 42:25–35.

18. Kurokawa C, Lynn GE, Pedra JHF, Pal U, Narasimhan S, Fikrig E. 2020. Interactions between *Borrelia burgdorferi* and ticks. Nat Rev Microbiol 18:587–600.

19. Zhu H, Fan J, Du J, Peng X. 2016. Fluorescent probes for sensing and imaging within specific cellular organelles. Acc Chem Res 49:2115–2126.

20. Xu W, Zeng Z, Jiang JH, Chang YT, Yuan L. 2016. Discerning the chemistry in individual organelles with small-Molecule fluorescent probes. Angew Chem Int Ed Engl 55:13658–13699.

21. Severo MS, Choy A, Stephens KD, Sakhon OS, Chen G, Chung DW, Le Roch KG, Blaha G, Pedra JH. 2013. The E3 ubiquitin ligase XIAP restricts *Anaplasma phagocytophilum* colonization of *Ixodes scapularis* ticks. J Infect Dis 208:1830–1840.

22. Jiang F, Doudna JA. 2017. CRISPR-Cas9 structures and mechanisms. Annu Rev Biophys 46:505–529.

23. Singh P, Schimenti JC, Bolcun-Filas E. 2014. A mouse geneticist’s practical guide to CRISPR applications. Genetics 199:1–15.

24. Kondo H, Rabouille C, Newman R, Levine TP, Pappin D, Freemont P, Warren G. 1997. p47 is a cofactor for p97-mediated membrane fusion. Nature 388:75–8.

25. Zhang S, Guha S, Volkert FC. 1995. The *Saccharomyces* SHP1 gene, which encodes a regulator of phosphoprotein phosphatase 1 with differential effects on glycogen metabolism, meiotic differentiation, and mitotic cell cycle progression. Mol Cell Biol 15:2037–50.

26. Kurtti TJ, Mattila JT, Herron MJ, Felsheim RF, Baldridge GD, Burkhardt NY, Blazar BR, Hackett PB, Meyer JM, Munderloh UG. 2008. Transgene expression and silencing in a tick cell line: A model system for functional tick genomics. Insect Biochem Mol Biol 38:963–8.

27. Chaverra-Rodriguez D, Macias VM, Hughes GL, Pujhari S, Suzuki Y, Peterson DR, Kim D, McKeand S, Rasgon JL. 2018. Targeted delivery of CRISPR-Cas9 ribonucleoprotein into arthropod ovaries for heritable germline gene editing. Nat Commun 9:3008.

28. Gilbert LA, Horlbeck MA, Adamson B, Villalta JE, Chen Y, Whitehead EH, Guimaraes C, Panning B, Ploegh HL, Bassik MC, Qi LS, Kampmann M, Weissman JS. 2014. Genome-scale CRISPR-mediated control of gene repression and activation. Cell 159:647–61.

29. Gilbert LA, Larson MH, Morsut L, Liu Z, Brar GA, Torres SE, Stern-Ginossar N, Brandman O, Whitehead EH, Doudna JA, Lim WA, Weissman JS, Qi LS. 2013. CRISPR-mediated modular RNA-guided regulation of transcription in eukaryotes. Cell 154:442–51.

30. Xia B, Viswanatha R, Hu Y, Mohr SE, Perrimon N. 2023. Pooled genome-wide CRISPR activation screening for rapamycin resistance genes in *Drosophila* cells. Elife 12:e85542.

31. Sajid A, Matias J, Arora G, Kurokawa C, DePonte K, Tang X, Lynn G, Wu MJ, Pal U, Strank NO, Pardi N, Narasimhan S, Weissman D, Fikrig E. 2021. mRNA vaccination induces tick resistance and prevents transmission of the Lyme disease agent. Sci Transl Med 13:eabj9827.

32. Commins SP. 2020. Diagnosis & management of α-gal syndrome: lessons from 2,500 patients. Expert Rev Clin Immunol 16:667–677.

33. Al-Rofaai A, Bell-Sakyi L. 2020. Tick cell lines in research on tick control. Front Physiol 11:152.

34. Carlyon JA. 2005. Laboratory maintenance of *Anaplasma phagocytophilum*. Curr Protoc Microbiol Chapter 3:Unit 3A 2.

35. Yoshiie K, Kim HY, Mott J, Rikihisa Y. 2000. Intracellular infection by the human granulocytic ehrlichiosis agent inhibits human neutrophil apoptosis. Infect Immun 68:1125–33.

36. Labandeira-Rey M, Skare JT. 2001. Decreased infectivity in *Borrelia burgdorferi* strain B31 is associated with loss of linear plasmid 25 or 28-1. Infect Immun 69:446–55.

37. Zuckert WR. 2007. Laboratory maintenance of *Borrelia burgdorferi*. Curr Protoc Microbiol Chapter 12:Unit 12C 1.

38. Abcam. 2022. Subcellular fractionation protocol.

39. Gasiunas G, Barrangou R, Horvath P, Siksnys V. 2012. Cas9-crRNA ribonucleoprotein complex mediates specific DNA cleavage for adaptive immunity in bacteria. Proc Natl Acad Sci U S A 109:E2579–86.

40. Ran FA, Hsu PD, Wright J, Agarwala V, Scott DA, Zhang F. 2013. Genome engineering using the CRISPR-Cas9 system. Nat Protoc 8:2281–2308.

41. Yang L, Yang JL, Byrne S, Pan J, Church GM. 2014. CRISPR/Cas9-directed genome editing of cultured cells. Curr Protoc Mol Biol 107:31.1.1-31.1.17.

42. Gantz VM, Bier E. 2015. Genome editing. The mutagenic chain reaction: a method for converting heterozygous to homozygous mutations. Science 348:442–4.

