## Supplementary Information for "Genetic manipulation of an *Ixodes scapularis* cell line"

**Supplementary Materials:**

Figures S1 to S2

Tables S1 to S2

### Supplementary Figure Legends

**Figure S1. Schematics of the ribonucleoprotein (RNP) delivery and selection in tick cells.** 200 pmol of the Cas9-NLS-tag RFP protein (Cas9 endonuclease tagged with nuclear localization sequence and red fluorescent protein) and 100 µM of sgRNA 1 and 2 were mixed and incubated at room temperature to prepare the RNPs. The RNP complex together with 7 µg of the donor DNA was nucleofected in 3 X $10^7$ ISE6 cells using the program code EN150 and the buffer SF on the 4D-Nucleofector System. ISE6 cells were then passaged for 7 cycles followed by puromycin selection, genomic DNA isolation and gene amplification analysis.

**Figure S2. Puromycin kill curve in tick cells.** 0.5 X $10^6$ ISE6 cells were seeded in a 12-well plate. After 24 hours, fresh L15C300 (Leibovitz-15 media) media with puromycin (0-10 µg/ml) was added to the cells. ISE6 cells were replenished with fresh media and antibiotic every 48 hours for 10 days. The viability of ISE6 cells was estimated using trypan blue.

**Supplementary Table 1**: Resources and reagents available

| **Antibody** | **Source** | **Identifier** | **Dilution/Concentration** |
|---|---|---|---|
| Mouse Anti-*I. scapularis* Relish monoclonal Ab | GenScript | custom | 1:500 |
| Histone H3 Rabbit polyclonal Ab | Cell Signaling | 9715 | 1:1,000 |
| GAPDH (14C10) Rabbit mAb | Cell Signaling | 2118S | 1:1,000 |
| Goat Anti-Rabbit IgG H&L (HRP) | Abcam | ab97051 | 1:4,000-10,000 |
| Rabbit Anti-Mouse Actin | Millipore Sigma | A2103 | 1:4,000 |
| HA-Tag (C29F4) Rabbit mAb | Cell Signaling | 3724 | 1:1,000 |
| Anti-Flag | Millipore Sigma | F3165 | 1:1,000 |
| Anti-NSFL1C mouse monoclonal Ab | Abnova | H00055968-B01P | 1:500 |
| Rat monoclonal [H139-52.1] Anti-Mouse kappa light chain (HRP) | Abcam | ab99632; discontinued | 1:4,000 |
| Rabbit Anti-*I. scapularis* XIAP polyclonal Ab | Thermo Scientific | custom | 1:1,000 |
| Rabbit Anti-*I. scapularis* Kenny polyclonal Ab | GenScript | custom | 1: 1,000 |
| Goat Anti-mouse IgG H+L(HRP) | Elabscience | E-AB-1001 | 1:3,000 |
| **Cell Dye** | | | |
| CellBrite™ Fix membrane stain | Biotium | 30090 | 1:1,000 |
| Hoechst 33342 Nuclear Stain | AAT Bioquest | 17533 | 1:1,000 |
| SlowFade Gold Antifade mountant | Invitrogen | S36938 | N/A |
| CytoPainter LysoGreen indicator reagent | Abcam | ab176826 | 1:1,000 |
| BioTracker™ 488 Green Mitochondria dye | Millipore | SCT136 | 1:1,000 |
| Cell Navigator™ TMR Ceramide Golgi staining kit- red fluorescence | AAT Bioquest | 22752 | |

| | | | |
|---|---|---|---|
| CytoPainter ER staining kit - green fluorescence | Abcam | ab139481 | 1:1,000 |
| **Restriction Enzymes** | | | |
| *SacI*-HF® | New England BioLabs | R3156S | N/A |
| *EcoRI*-HF® | New England BioLabs | R3101S | N/A |
| *BamHI*-HF® | New England BioLabs | R3136S | N/A |
| *SalI*-HF® | New England BioLabs | R3138S | N/A |
| **Cell Media** | | | |
| Leibovitz's L-15 Medium, powder | Gibco | 41300039 | N/A |
| L-aspartic acid | Millipore-Sigma | 11189 | 0.449 g/L |
| L-glutamine | Millipore-Sigma | G8540 | 0.500 g/L |
| L-proline | Millipore-Sigma | 81709 | 0.450 g/L |
| L-glutamic acid | Millipore-Sigma | 49449 | 0.250 g/L |
| α-ketoglutaric acid | Millipore-Sigma | K1128 | 0.449 g/L |
| Sodium hydroxide | Millipore-Sigma | S8045 | 10 N |
| D-glucose | Millipore-Sigma | G7021 | 18.018 g/L |
| FBS (USDA approved; for tick medium) | Millipore-Sigma | F0926-500ML | 10% |
| Bacto™ Tryptose Phosphate Broth | BD | 260300 | 10% |
| Lipoprotein concentrate | MP Biomedicals | 191476 | 0.10% |
| Normal rabbit serum | Pel-Freez | #31126-5 | 6.00% |
| Sodium bicarbonate | Millipore-Sigma | S6014 | 0.25% |
| HEPES | Millipore-Sigma | H4034 | 25 mM |
| CMRL1066 w/L-Glutamine (Powder) | US Biological | C5900 | N/A |
| Sodium citrate tribasic dihydrate | Millipore-Sigma | S4641 | 0.7 g/L |
| Yeastolate | BD | 255772 | 2 g/L |
| Neopeptone | BD | 211681 | 5 g/L |
| N-Acetyl-α-D-glucosamine | Millipore-Sigma | 1079-25GM | N/A |

| | | | |
|---|---|---|---|
| Albumin, Bovine Fraction V | MP Biomedicals | 160069 | N/A |
| Rifampicin | Millipore-Sigma | 557303 | 50 ug/ml |
| Phosphomycin | Millipore-Sigma | P5396 | 100 ug/ml |
| Amphotericin B | Gibco | 15290-026 | 0.111111111 |
| Sodium pyruvate | Millipore-Sigma | P5280 | 0.8 g/L |
| RPMI-1640 Medium With L-Glutamine | Quality Biological | 112-025-101 | N/A |
| Fetal Bovine Serum (for HL-60s media) | Gemini Bio-Products | 100-106 | 0.1 |
| 1X PBS | Quality Biological | 114-058-101 | N/A |
| Distilled water | Gibco | 15-230-147 | N/A |
| LB broth | Millipore-Sigma | L3022 | N/A |
| LB agar with 100 ug/ml ampicillin | Quality Biological | 50-751-7582 | N/A |
| Ampicillin | Millipore-Sigma | A0166 | 100 ug/ml |
| **Reagents and Materials** | | | |
| Cellstar® cell culture flasks, 25 $cm^2$ | Greiner bio-one | 690-160 | N/A |
| T-25 Vented Flasks | CytoOne | CC7682-4825 | N/A |
| Cell culture plate with lid (6 well, flat bottom) | Millipore-Sigma | SIAL0516 | N/A |
| iTaq™ Universal SYBR® Green Supermix | Biorad | 1725121 | N/A |
| Costar® cell culture plate with lid (24 well, flat bottom) | Corning | CLS3526-1EA | N/A |
| Mini-Protean® TGX™ gels | Biorad | 456-9034 | N/A |
| Trans-blot® Turbo™ Transfer pack, 0.2 µm PVDF | Biorad | 1704156 | N/A |
| Miller GP 0.2 µm filter unit | Millipore-Sigma | SLGP033RS | N/A |
| FALCON® 14 ml polypropylene round-bottom tube | Corning | 352059 | N/A |
| Nunc™ 96-Well polystyrene round bottom microwell plates | Thermo Scientific | 262162 | N/A |
| 250 mm glass desiccator | Fisher Scientific | 08-615B | N/A |
| 1.5 ml microcentrifuge tubes | Thomas Scientific | 1148T71 | N/A |

| | | | |
|---|---|---|---|
| 15 ml conical screw cap tubes | USA Scientific | 5618-8261 | N/A |
| 50 ml conical screw cap tubes | USA Scientific | 5622-7270 | N/A |
| 500 ml vacuum filter/storage bottle system, 0.2 μm | Corning | 430773 | N/A |
| 27 gauge, 1/2" needle | BD | 305109 | N/A |
| White filter paper for CytoSep™ single funnel | Simport Scientific | M965FW | N/A |
| Richard-Allan Scientific™ three-step stain set | Thermo Scientific | 3300 | N/A |
| Epredia™ Cytospin™ 4 cytocentrifuge | Thermo Scientific | A78300003 | N/A |
| Fisherbrand™ Superfrost™ plus microscope slides | Thermo Scientific | 12-550-15 | N/A |
| Halt™ phosphatase inhibitor cocktail (100x) | Thermo Scientific | 78426 | 1:100 |
| Halt™ protease inhibitor cocktail (100x) | Thermo Scientific | 87786 | 1:100 |
| 10X RIPA | Millipore-Sigma | 20-188 | 1X |
| Chloroform | Millipore-Sigma | 288306 | 100% |
| TRIzol reagent | Ambion | 15596018 | N/A |
| Methanol anhydrous, 99.8% | Millipore-Sigma | 322415-1L | 100% |
| Ethyl alcohol, pure; 200 proof for molecular biology | Millipore-Sigma | E7023-1L | 70-100% |
| HyClone™ water, molecular biology grade | Cytiva | SH3053801 | N/A |
| Puromycin dihydrochloride | Millipore-Sigma | P9620 | 4 μg/ml |
| Pierce™ Anti-dykddddk affinity resin | Thermo Scientific | A36803 | N/A |
| Pierce™ IP lysis buffer | Thermo Scientific | 87787 | N/A |
| 2-β mercaptoethanol | Gibco | 21985-023 | 5% |
| 6X Laemmli buffer | Alfa Aesar | J60660 | 1X |
| Blocking grade blocker, non-fat skim milk | Biorad | 1706404 | 5% |
| Bovine serum albumin | Millipore-Sigma | A2058 | 3% |
| Sodium dodecyl sulfate (SDS) | Millipore-Sigma | L6026 | 0.10% |

| | | | |
|---|---|---|---|
| Paraformaldehyde | Millipore-Sigma | P6148 | 4.00% |
| JumpStart™ REDTaq® ReadyMix™ Reaction Mix | Millipore Sigma | P1107 | 1X |
| Phusion® HF DNA Polymerase | New England BioLabs | M0530S | 1 unit |
| **Commercial Assays** | | | |
| Pure Link RNA mini kit | Ambion | 12183025 | N/A |
| SF Cell Line 4D-Nucleofector™ X Kit L | Lonza Bioscience | V4XC-2012 | N/A |
| Verso cDNA Synthesis Kit | Thermo Scientific | AB-1453B | N/A |
| Pierce BCA Protein Assay Kit | Thermo Scientific | 23227 | N/A |
| Pierce ECL Western Blotting Substrate | Thermo Scientific | 32106 | N/A |
| DNeasy® Blood and Tissue kit | Qiagen | 69506 | N/A |
| QIAamp® DNA Mini Kit | Qiagen | 51304 | N/A |
| QIAprep® Spin Miniprep Kit | Qiagen | 27106 | N/A |
| *Mycoplasma* testing kit | Southern Biotech | 13100-01 | N/A |
| Endofree Plasmid Maxi Kit | Qiagen | QGN-12362 | N/A |
| Immobilon Forte Western HRP substrate | Millipore-Sigma | WBLUF0100 | N/A |
| **Equipment** | | | |
| 4D-Nucleofector™ System | Lonza Bioscience | AAF-1002 | N/A |
| Cytospin 4 | Thermo Scientific | A78300003 | N/A |
| CFX96 Touch Real-Time PCR Detection System | Biorad | Discontinued | N/A |
| C1000 Touch Thermocycler | Biorad | 1851148 | N/A |
| **Cell Lines** | | | |
| *I. scapularis* ISE6 cells | Ulrike Munderloh, University of Minnesota | | ISE6 |
| **Organisms** | | | |
| One Shot Top10 Chemically Competent E. Coli | Thermo Scientific | C404003 | N/A |
| HL-60 cells | ATCC | CCL-240 | NA |
| *A. phagocytophilum* HZ strain | Ulrike Munderloh, University of Minnesota | | N/A |

| | | | |
|---|---|---|---|
| *B. burgdoferi* B31 clone MSK5 | Jon Skare, Texas A&M University Health Science Center | | N/A |
| **Plasmids** | | | |
| pAQUA-N1 | Addgene | 42888 | N/A |
| pCMV-HA | Addgene | 32530 | N/A |
| pCMV-FLAG | Sino Biological, Inc | CV002 | N/A |
| DsRed-N1 | Addgene | 54493 | N/A |
| Donor vectors | Origene Technologies | custom | N/A |
| **Protein** | | | |
| Cas9-NLS-tagRFP protein | Genaxxon bioscience | S5306.0010 | N/A |

**Supplementary Table 2: Primers used**

| Name | Type | Target Accession | Start position within mRNA or exon | Strand | Sequences (5'-3') |
|---|---|---|---|---|---|
| | | | 1 | F | GATTGCAG**GAATTC**ATGGTTGTCATCAGTATGGCG |
| **pCMV-*xiap*-HA*** | Cloning | XM_002433822 | 1014 | R | GTATAGTT**GCGGCCGC**TCATGAAAGAAAAGCCTTAATGTTC |
| | | | 27 | F | GGCC**GAATTC**ATGGCAGATTGTGCGGGGCG |
| **pCMV-*p47*-FLAG*** | Cloning | XM_002433590 | 1175 | R | GGCC**GTCGAC**TCACTTGATACGCTGGACGATAA |
| | | | 1 | F | ACTCAGATCTC**GAGCTC**GATGGTTGTCATCAGTATGGCGATC |
| ***xiap*-Dsred2-N1** | Cloning | XM_002433822 | 1014 | R | CGTCGACTGCA**GAATTC**CTGAAAGAAAAGCCTTAATGTTCTC |
| | | | 27 | F | ATTCTGCA**GTCGAC**GGATGGCGGACTGCGAGGAAC |
| ***p47*-AquaN1** | Cloning | XM_002433590 | 1175 | R | GCGACCGGT**GGATCC**CCTTTTATACGCTGCACAATG |
| | | | | LHA-F | TGGTAGCTAAAGGCCTACAC |
| ***p47*** | PCR | XM_002433590 | NA | RHA-R | GTCTTCACAAAGGGCAAAGG |
| | | | | LHA-F | TGGCGGTACCTTCCCAAATC |
| ***xiap*** | PCR | XM_002433822 | NA | RHA-R | AGTGTGCAACGGGATGCTAC |
| | | | | Puro-F | GCCTCTGTTCCACATACACTTC |
| **donor cassette** | PCR | NA | NA | RFP-R | GATGCCCTGGGTGTGGTTGATG |
| ***p47* sgRNA** | sgRNA | LOC120845803 | 498 | F | CTGTACGGAGAACGACACAG |

| | | | | | |
|---|---|---|---|---|---|
| | | | 468 | R | AGCAGCGGTCATCGCGCAAC |
| ***xiap* sgRNA** | sgRNA | LOC8050569 | 259 | F | AAGGCGGGACTTTTTTACAA |
| | | | 228 | R | TCGGCGACGAGTACGTACAG |
| ***A. phagocytophilum* 16S rRNA** | qRT-PCR | NC_007797 | 283 | Ap16S_F | CAGCCACACTGGAACTGAGA |
| | | | 400 | Ap16S_R | CCCTAAGGCCTTCCTCACTC |
| ***I. scapularis p47*** | qRT-PCR | XM_040217354.2 | 93 | p47_F | GCCAGGGCCAAGCTTTACC |
| | | | 190 | p47_R | CTTGGACGCTCCAGCGAC |
| ***I. scapularis kenny*** | qRT-PCR | XM_040500911.2 | 1169 | Key_F | GCTCAGGACTTGGCAGGAAT |
| | | | 1300 | Key_R | CACCAGCTTGTCTTGGACCT |
| ***I. scapularis relish*** | qRT-PCR | XM_040501061.2 | 487 | Relish_F | AGAATGTCCGCCACCGTTTTTTTCTGC |
| | | | 595 | Relish_R | CACGTGCACCGCCTCACCATGAAGG |
| ***I. scapularis actin*** | qRT-PCR | XM_029977298 | 896 | I.s. actin_F | GGTCATCACAATCGGCAAC |
| | | | 1003 | I.s. actin_R | ATGGAGTTGTACGTGGTCTC |

*Constructs used in McClure Carroll *et al.*, 2019

Bold letters are introduced restriction endo

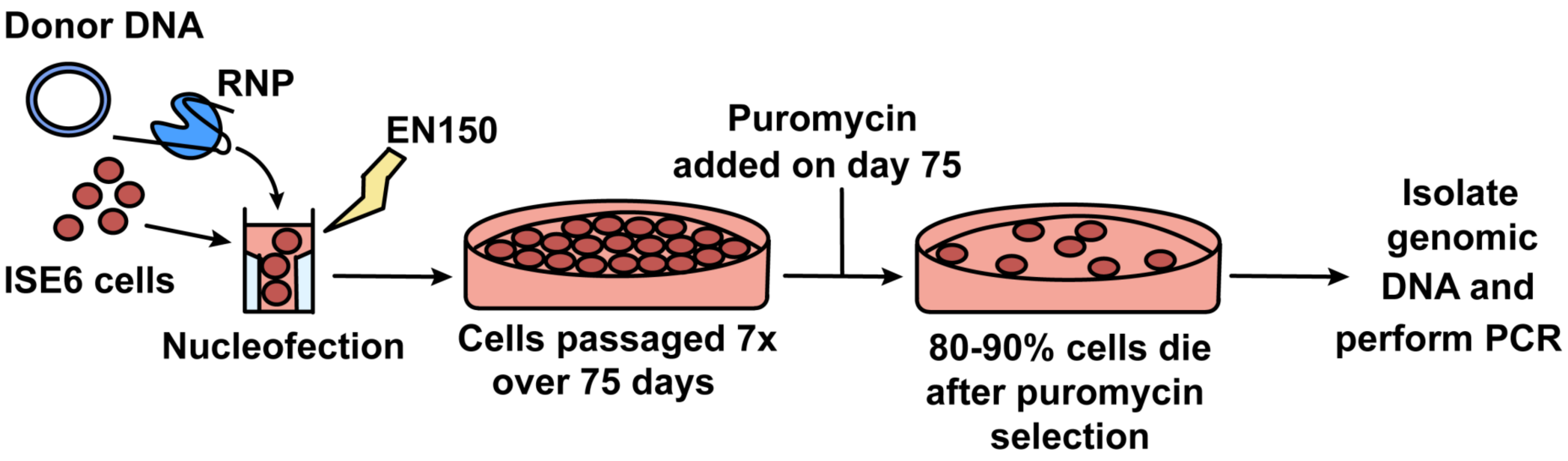


**Singh *et al.* Figure S1**

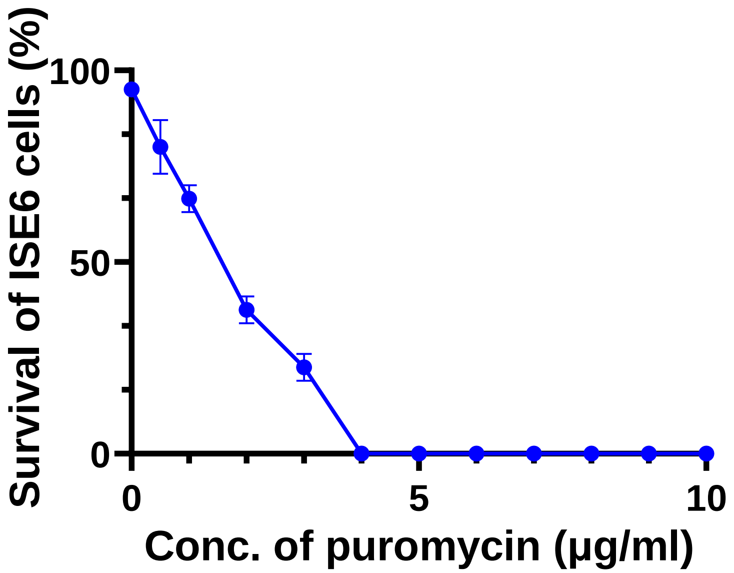


Singh *et al.* Figure S2
